# Degradation of white birch shelterbelts by attack of white-spotted longicorn beetle in central Hokkaido, northern Japan

**DOI:** 10.1101/2020.01.30.926188

**Authors:** Kazuhiko Masaka, Yohichi Wakita, Kenta Iwasaki, Masato Hayamizu

## Abstract

Widespread decline of white birch shelterbelts was observed in central Hokkaido, northern Japan. Many exit holes bored by adults of the white-spotted longicorn beetle have been found at the bases of the trunks of trees in these stands. The number of adult longicorn beetle exit holes (*N*_holes_) of dead standing trees tended to be greater than that of living trees. *N*_holes_ tended to increase with increasing *DBH*, and there was a negative relationship between *N*_holes_ and tree vigor. We found a size-dependent lethal threshold in *N*_holes_. A resonance-measurement device (RMD) for diagnosing the level of wood defection inside the trunk was also tested. The RMD examination together with the lethal threshold in *N*_holes_ can be a useful tool for the diagnosis of white birch trees. We estimated *N*_holes_ of dead standing trees with a *DBH* of 25 cm in each plot (*N*_D25_) to compare the severity of infestation among plots. Logistic regression analysis revealed that 50% of stands will be degraded if *N*_D25_ = 25.0. Thus, the degradation could also be evaluated by *N*_holes_.

## Introduction

The genus *Anoplophora* (Coleoptera) includes some of the most damaging wood-boring pests in the Northern Hemisphere (Haack et al. 2010; Hu et al. 2009; Meng et al. 2015; van der Gaag and Loomans 2014). Though *Anoplophora* spp. has significant preferences for particular host trees (Faccoli and Favaro 2016; Iwaizumi et al. 2014), it has a very wide host plant range that includes more than 100 tree species (Sjörman et al. 2014; also see van der Gaag and Loomans 2014). Larvae of *Anoplophora* spp. develop in the sapwood of living trees, and damage may kill the trees (Haack et al. 2010; Meng et al. 2015). However, we have limited information about the process of forest decline from the viewpoint of forest management (cf. Dodds and Orwig 2011), whereas many researchers have focused on the ecology of *Anoplophora* spp. from the viewpoints of dispersal behavior (Bancroft and Smith 2005; Favaro et al. 2015; Hull-Sanders et al. 2017; Smith et al. 2004; Williams et al. 2004), host preference (Faccoli and Favaro 2016; Fujiwara-Tsujii et al. 2016; Yasui and Fujiwara-Tsujii 2013) and invasion process (Javal et al. 2019a,b; Tsykun et al. 2019).

White-spotted longicorn beetle (*A. malasiaca*) native to Japan has long been notorious pest of fruit trees such as citrus, pear etc. and ornamental trees such as sycamore, maple etc. (Kojima and Nakamura 2011). In silviculture, damage of sugi cedar (*Cryptomeria japonica*: Taxodiaceae) plantations by this beetle has been also sometimes reported in western Japan (Kobayashi and Okuda 1981; Taniguchi et al. 1982). Recently, degradation of shelterbelts composed of Japanese white birch (*Betula platyphylla* var. *japonica*: Betulaceae), has been observed in central Hokkaido, Japan (Masaka 2017; Fig. 1A). Abundant holes with a diameter of ca. 1 cm were often found along the basal region of the trunks (up to ca. 50 cm above ground; Fig. 1B) and exposed roots (Fig. 1C), even in apparently intact shelterbelts (Masaka 2017). Some holes were clogged by fresh frass. In Hokkaido, four major wood-boring pests; *A. malasiaca*, *Buprestidde agrilus* (Buprestidae), *Opostegoides minodensis* (Opostegidae), and *Phytobia betulae* (Agromyzidae), that attack the living birch wood were reported by Hara (2000), and the features mentioned above indicated infestation of white-spotted longicorn beetle (Hara Hideho, personal communication). Actually, we often found the adult white-spotted longicorn beetles in infested stands (Masaka 2017), and the larvae were also found in the wood (Fig. 1D). However, birch species is unlikely to be a major favorite host for *Anoplophora* spp. compared with citrus, maple and popular (cf. Iwaizumi et al. 2014; Sjörman et al. 2014; van der Gaag and Loomans 2014), and only one case of serious infestation of white birch (*B. platyphylla*) by Asian longhorned beetle (*A. glabripennis*) was reported in China (Gao et al. 2009). In Japan, mass attack on Japanese white birch was sporadically reported at the stand level, but it did not cause to death (Makihara et al. 1989; Onodera et al. 1995, 1997). Thus massive mortality of Japanese white birch has not ever been reported in Japan.

**Figure 1.**
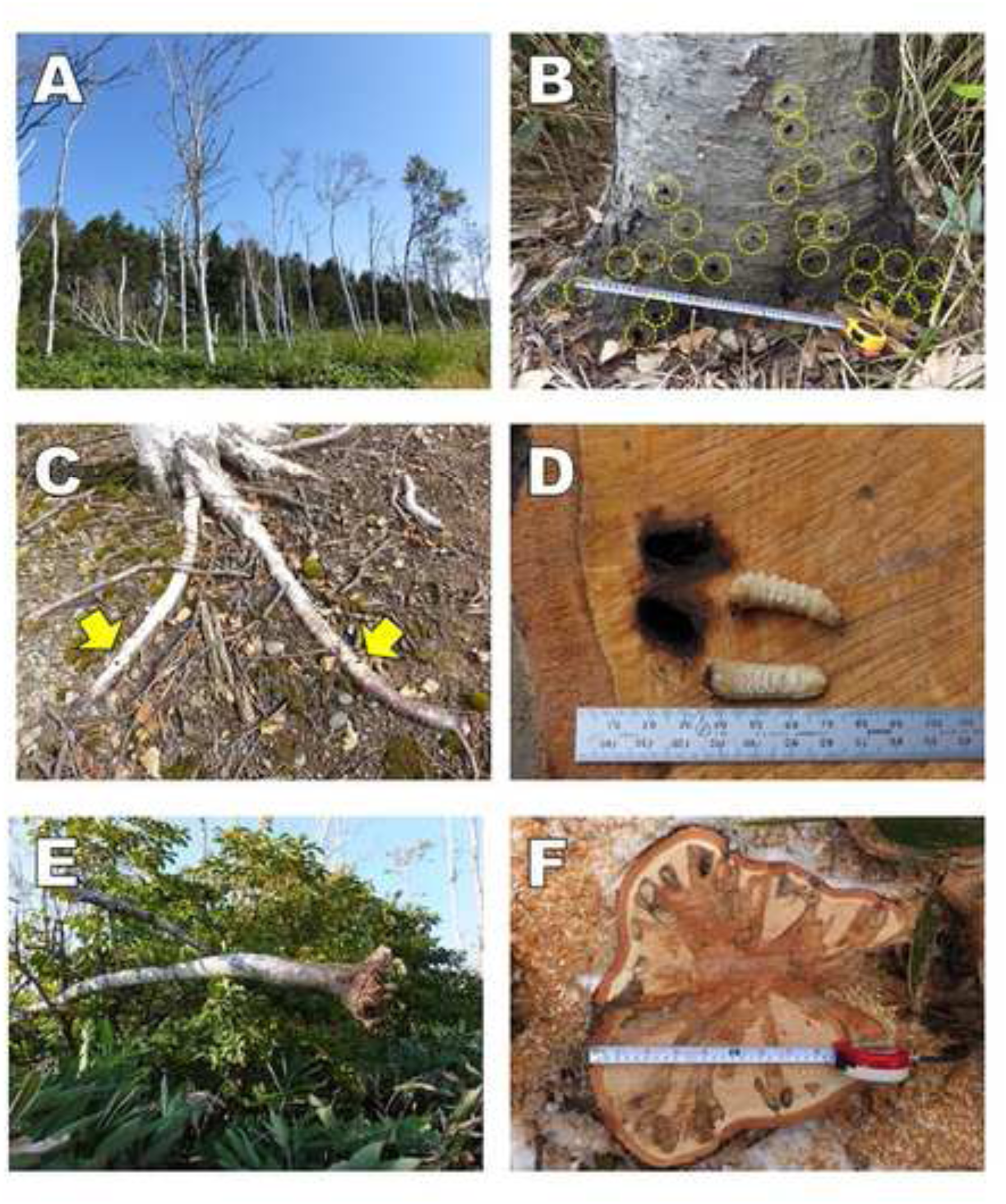
Infestation of Japanese white birch shelterbelts by white-spotted longicorn beetle. Following location and plot number are shown in Fig. 2 and Table S1. A) Degraded stand at Plot 7 in Bibai (21 Sep., 2016). B) Adult exit holes observed at the trunk base (dotted circles) at Plot 15 near the degraded stands; i.e., Plot 12 and Plot 13. Though this stand seemed to be intact in appearance, 1/3 of standing trees were dead (Table S1; 25 Sep., 2015). C) Adult exit holes on the root as shown by the arrows (14 Oct., 2016; In Shinshinotsu, from Masaka [2017]). D) Larvae of white-spotted longicorn beetle in the sapwood (5 Dec., 2016). E) Snapping of trees at the trunk base found at Plot 13 (26 Sep., 2016; from Masaka [2017]). F) Cross section of wood at ground level. Palmate discolored area elongates toward the tunnel bored by the larvae (1 Dec., 2016).

Dead standing trees of white birch are often observed in the old-growth stands (cf. Takahashi et al. 1974). The dead trees lose their shoots, twigs, and eventually large branches in the crown, so that eventually only the trunks remain, which will then decay over time. However, many fallen trees have also been found in the degraded stands, some of which still had crowns with leaves still on the twigs (Masaka 2017). These fallen trees had snapped at the trunk base without being uprooted (Fig. 1E); this indicates that the trunk wood has been severely damaged. In some cases, white-spotted longicorn beetle larvae had tunneled into the wood so much that it caused a ‘swiss cheese’ appearance of the wood (Masaka 2017; Fig. 1F). It will cause the mechanical vulnerability of the root system. Falling trees may injure passers-by on the farm road along the shelterbelts, and fallen logs on farmland may pose significant issues for farmers. As there are many white birch shelterbelts in central Hokkaido (cf. Sato et al. 2009), management decisions must be made quickly about whether the stand should be removed immediately or not. Therefore, we have to assess the extent of the damage of white birch shelterbelts caused by the longicorn beetle in central Hokkaido and demonstrate the risk of fallen trees with respect to the severity of infestation. From the view point of forest management rather than the pest control, we aimed to investigate whether the number of adult exit holes can be used as an index of the decline of trees in shelterbelts. In addition to the visual inspection, we also applied the decay diagnosing method of wood using the resonance measurement device (RMD) in corroboration of the effect of infestation on the wood.

In the present study, we aimed to address four questions: (1) To what extent does the number of adult exit holes cause the decline of white birch? (2) Is there any threshold for the mortality of individual trees in relation to the severity of infestation? (3) To what extent can RMD be used to measure the wood condition of infested trees? (4) To what extent does the severity of infestation cause the degradation at the stand level?

We have used *A. malasiaca* as the scientific name in this study to follow the Dictionary of Japanese Insect Names (http://konchudb.agr.agr.kyushu-u.ac.jp/dji/), though *A. malasiaca* is often used as a synonym of *A. chinensis* (Haack et al. 2010). DNA analysis has revealed that *A. malasiaca* is not necessarily the same as *A. chinensis* in Japan (Muraji et al. 2011).

## Materials and methods

### Study area

We carried out the investigation in Bibai, Mikasa, Iwamizawa, Tsukigata and Shinshinotsu in central Hokkaido (Fig. 2). Shelterbelts in central Hokkaido were established after the Pacific War on drained wetlands around the Ishikari River system (Fig. 2 and Table S1; IBCH 1970). In this region under cool and humid conditions, main purpose of establishing the shelterbelts and windbreaks was the promotion of crop growth due to increase in temperature (cf. Iwasaki et al. 2019). In addition to white birch, Manchurian ash (*Fraxinus mandshurica* var. *japonica*: Oleaceae), Dahurian larch (*Larix dahurica* var. *japonica*: Pinaceae) and Norway spruce (*Picea abies*: Pinaceae) have been used in the establishment of shelterbelts (cf. Sato et al. 2009). Each shelterbelt is composed of either a single species or a combination of species arranged in a line (Fig. 1A). Each shelterbelt has a length of ca. 500–3000 m and a width of ca. 20–50 m. The ground vegetation is often dominated by dwarf bamboo (*Sasa senanensis*: Poaceae). Narrower ‘windbreaks’ that consist of a few rows of trees (ca. 2–3 m in width) are also occasionally found in this region. The shelterbelts were established by the municipal or national forestry administrations, whereas the windbreaks were established by private organizations or individuals.

**Figure 2.**
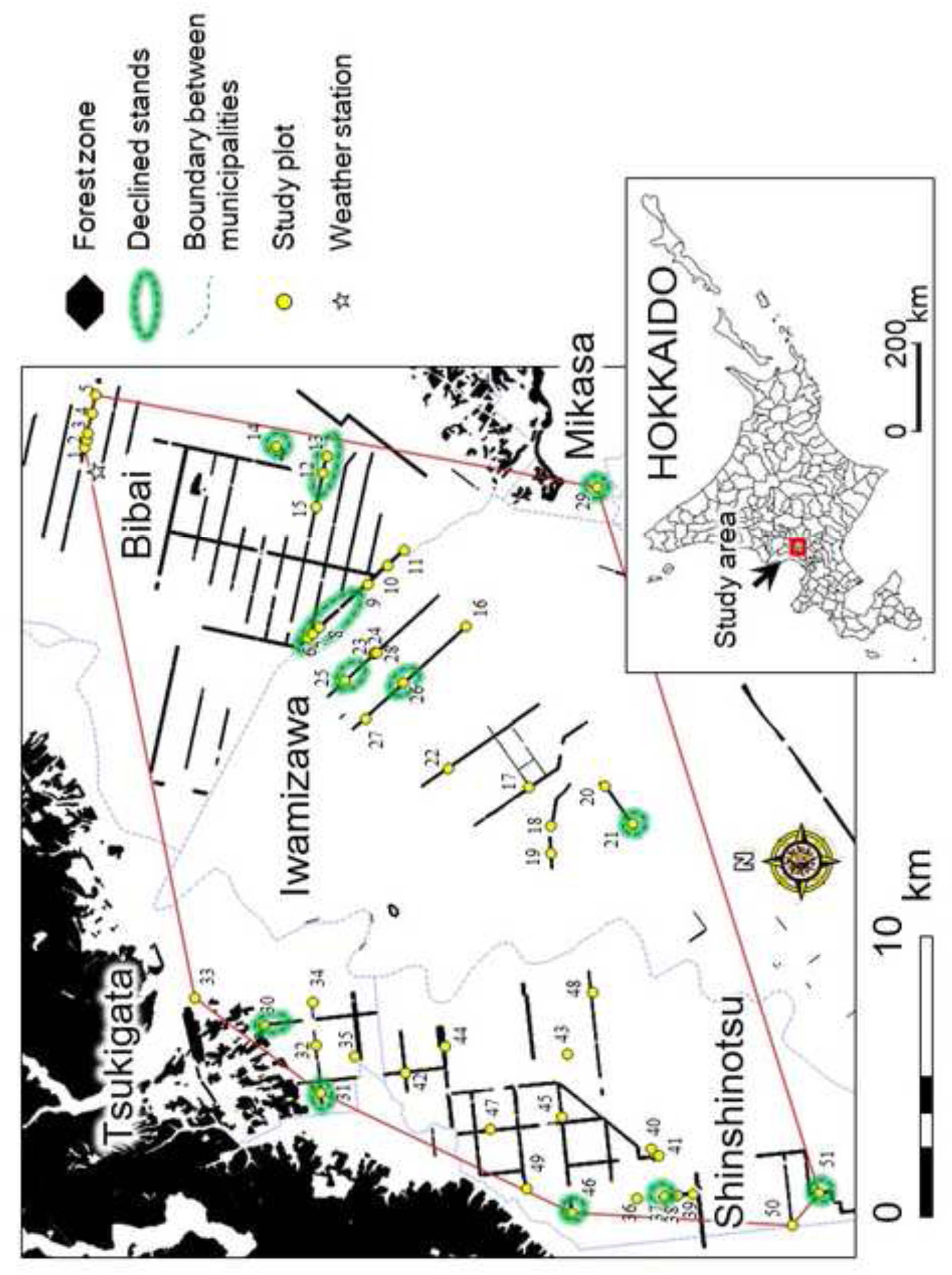
Study area. Polygon indicates study area.

According to the Japan Meteorological Agency (1981–2010; http://www.data.kishou.go.jp/), annual precipitation in Bibai (see Fig. 2) is 1156.5 mm year^−1^, and the mean monthly temperatures during the warmest (August) and coldest (January) months are 26.4 °C and −12.0 °C, respectively. There is snow cover during November–April, with a mean annual maximum depth of 116 cm.

### Relationship between the number of adult exit holes of the longicorn beetle and tree vigor

We propose two assumptions for the evaluation of the effect of infestation. First, the maximum cumulative capacity of each tree to hold the larvae that can pupate and develop into adults is size dependent. It means that there is an upper limit in wood volume as a food resource and living space for the larvae. Then adult emergence in each tree will increase with increasing tree size. Second, decline of tree is affected by the severity of infestation with respect to the tree size. From these viewpoints, we can expect that the relationship between number of adult exit holes and trees size differs according to the tree vigor. Besides, there would be a lethal threshold in the number of adult exit holes with respect to the tree size.

In August–October 2015, we established 15 study plots in shelterbelts in Bibai (Figs. 1A and 2, and Table S1). Six plots out of the 15 plots were obviously in degraded stands (Plot 6, 7, 8, 12, 13, 14; Fig. 1A and Table S1). We defined the degraded stands as the number of dead standing trees > 30% in the stand, but some stands were entirely destroyed (Fig. 1A and Table S1). We counted the number of adult exit holes (*N*_holes_) and measured the diameter at breast height (*DBH*; cm) of all living and dead standing trees in each plot. The crown vigor of each tree in the plots (hereafter *Vigor*; no unit) was assessed according to the following scale (cf. Masaka et al. 2000): (0) no foliage in the crown, only a standing trunk remained, which was considered to be dead (dead standing tree); (1) less than 1/10 of the foliage remaining in the crown; (2) 1/10–1/4 of the foliage remaining; (3) 1/4–1/2 of the foliage remaining; (4) 1/2–3/4 of the foliage remaining; (5) more than 3/4 of the foliage remaining (intact crown). To assess the combination between tree vigor and *N*_holes_–*DBH* relationship, we conducted a generalized linear mixed model (GLMM) analysis as follows:

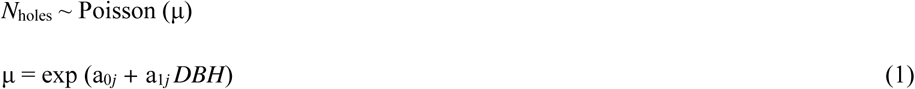

This is the base model at the individual level. The model assumed that both the intercept (a_0*j*_) and slope (a_1*j*_) of *N*_holes_–*DBH* relationship in *j*-th plot are linked with degree of tree vigor (*Vigor*) as follows:

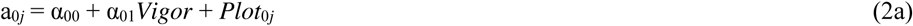

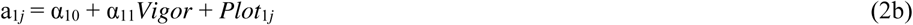

where a*_ij_* and α*_ik_* (*i* = 0, 1; *k* = 0, 1) are regression coefficients. Thus the model represent the *N*_holes_–*DBH* relationship with different *Vigor*s. *Plot_ij_* (*i* = 0, 1) is the random effect specific to each plot, in which *Plot*_0*j*_ indicates the random intercept and *Plot*_1*j*_ indicates the random slope at the *j*-th plot. Though *Vigor* is the ordinal variable, we treat it as the numerical variable to simplify the interpretation of the result. A Poisson distribution was assumed for the objective variable.

### Use of a resonance-measurement device to diagnose intact stands

We trialed the use of a resonance-measurement device (RMD; Tree Health Checker, Ebisu System Co.,Ltd., Sapporo) to diagnose the degree of wood decay. This device was developed jointly by Hokkaido Research Organization and Ebisu System Co.,Ltd. The RMD diagnoses the level of wood defection inside a tree trunk based on the principle that homogeneous solid materials show a narrow range of elastic wave velocities, which can be measured by the resonance of the material (Japanese patent application No. 5531251). Signals were applied by a shaker to the surface of the trunk and a receiver was placed on the opposite side of the trunk. Measurements were made at 12 positions around the girth of each trunk at 90° intervals. Defects in the wood cause great deviation among the measurements. Therefore, diagnosis was based on the following tests (threshold values are patent pending): (1) deviation of the resonance frequency in both a fundamental wave and a harmonic wave; (2) deviation of the resonance frequency among measurements from different positions; and (3) divergence between the species-specific resonance frequency and the resonance frequency of the focal trees. Threshold values are not described in this paper, as the information is subject to a pending patent application. If a focal tree did not pass any of these tests, its wood was judged as *critical*, as the inside of the tree trunk was evidently seriously rotten. Diagnoses of *critical* and *suspicious* (requires further examination) were based on patent-pending values. *Not bad* definitely indicated that there was no wood defect.

We conducted this analysis on two stands in Bibai (Plot 9 and Plot 15; see Table S1). These stands did not show obvious signs of degradation apparently, but there were many adult exit holes at the base of the trunks. We used the RMD to measure (on 14 Apr., 2017) all living trees (n = 59) and two dead trees that had been alive in the previous year at these sites. Measurements were made at a height of 0.5 m above the ground, since the measurement requires cylindrical trunk for the conduction of elastic wave. The round cross section of trunk often loses its shape near the ground surface because of buttress (cf. Fig. S1). Prior to this investigation, we had measured the wood of 12 white birches in other stands (including two trees that had not been infested by the longicorn beetle) on 21 Sep., 2016. These trees were then cut on 1 Dec., 2016 and the degree of discoloration in the cross-section of the wood was assessed to adjust the calibration (Fig. S1).

If the severity of infestation would reflect on the diagnosis, we could expect that *N*_holes_–*DBH* relationship also differs according to diagnoses. To evaluate the connection between *Diagnosis* and *N*_holes_‒ *DBH* relationship with attention to the random effect, we used the base model written in Eq. 1. Similar to Eqs. 2a and 2b, the model assumed that both the intercept (a_0*j*_) and slope (a_1*j*_) of *N*_holes_–*DBH* relationship are linked with *Diagnosis*, with attention to the plot-specific character as random effect (*Plot_ij_*; *i* = 0, 1) as follows:

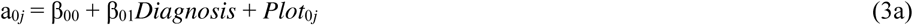

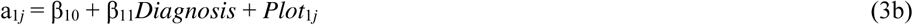

where β*_ik_* (*i* = 0, 1; *k* = 0, 1) is the regression coefficient. Thus the model represents the *N*_holes_–*DBH* relationship with different *Diagnosis*s. *Diagnosis* is a categorical variable with values of *not bad*, *suspicious,* and *critical*.

### Probability of the appearance of degraded stands with respect to the severity of infestation

Degradation of the stands was evaluated by *N*_holes_ in dead standing trees. *N*_holes_ does not increase after the death of the host tree, since the longicorn beetle only infests living trees. Then the statistics of *N*_holes_ in dead trees will be the representative index specific to each plot, i.e., it indicates the severity of infestation in each plot.

In addition to the 15 plots in Bibai outlined above, we established a further 36 plots in shelterbelts (*n* = 30), windbreaks (*n* = 2), trees lining road-sides (*n* = 2), and natural forests (*n* = 2) around Bibai (Iwamizawa, Mikasa, Shinshinotsu, and Tsukigata) from October 2016 to June 2017 (Figs. 2 and S2). These plots included three stands composed of Erman’s birch (*B. ermanii*) and two of silver birch (*B. verrucosa*) (Table S1). We selected all severely degraded stands as long as possible (Plot 21, 24, 25, 26, 29-31, 37, 46, 51; Fig. S2), and then, selected apparently intact stands arbitrarily across the region. Notes that almost all dead trees were removed in Plot 30 and Plot 51 in forest practice before investigation. Besides, there were many trees with a little foliage in the crown in Plot 21 (personal observation) presumably because of many *N*_holes_ (see Table S1). We considered that the trees in Plot 21 would be just before dead. On the other hand, four stands were intact in appearance (Plot 9, 32, 38, 41), many adult exit holes were observed at the trunk base and many dead canopy trees were observed (see Table S1). We considered that these four stands reached a stage just before degradation, but these were included in the analysis as intact stands. Similar to the previous investigation in Bibai, we counted *N*_holes_ and measured the *DBH* of all living and dead standing trees in each plot.

Probability of the appearance of the degraded stands with respect to *N*_holes_ was evaluated by the logistic regression analysis (degraded stand = 1, intact stand = 0). As the tree size distribution differed among plots according to the different stand ages and site qualities, we compared the severity of infestation among plots for same-sized trees. Therefore, we estimated the average *N*_holes_ for dead standing trees with a *DBH* of 25 cm (hereafter referred to as *N*_D25_) in each plot using Eq. 1. *N*_D25_ of dead standing trees in each plot was obtained by substituting *DBH =* 25 cm in the model. In the logistic regression analysis, binomial distribution was assumed for the objective variable.

### Statistical analysis

We used R ver. 3.4.4 (2018 The R Foundation for Statistical Computing) for GLMM analyses, with the *glmer* function in the package *lme4*. Generalized linear model (GLM) was conducted instead of GLMM, if there was strong correlation between the random effects. Logistic regression analysis was conducted using *glm* function with logit link. Coefficients of the random intercept and random slope for each plot were output by the *ranef* function. To assess the best combination of the fixed variables, package *MuMIN* was used for model selection. Akaike’s information criterion (AIC), which balances the fit of the model against the number of parameters, was used to select the best-fit model for the GLMM (Burnham and Anderson 2002). The model with the smallest AIC value was accepted as the best fit for the data (Crawley 2005; McCarthy 2007). The model selection in Eq. 3 was conducted to pay attention to the combination of the categories in *Diagnosis* (*not bad* [*N*], *suspicious* [*S*], *critical* [*C*]) – i.e., five combinations of the categories can be defined: (1) the three categories differ from each other (*N ≠ S ≠ C*); (2) *S* = *C ≠ N*; (3) *N* = *S ≠ C*; (4) *N* = *C ≠ S*; (5) the three categories did not differ from each other (*N* = *S* = *C*).

## Results

### Relationship between the decline of white birch and infestation by the longicorn beetle

In our examination of wood cross-sections, we often observed palmate discolored areas, each finger of which appeared to extend toward a tunnel at the sapwood (Figs. 1D and F; cf. EPPO, 2007). Furthermore, the tunnels were often clogged by rotten and blackening sawdust (Fig. S3), which could be a source of infection by wood rot fungi (cf. Figs. 1F and S2) and eventually lead to the snapping of trees at the trunk base (Fig. 1E).

Frequency distribution of *N*_holes_ differed markedly among tree vigors, and *N*_holes_ of dead standing trees (*Vigor* = 0) was greater than that of living trees (*Vigor* = 1 – 5) (Fig. 3A). The *N*_holes_–*DBH* relationship for living and dead white birch is shown in Fig. 3B. As expected, *N*_holes_ increased with increasing *DBH*. However, the nature of this relationship appeared to differ between living and dead trees. In the GLMM analysis, both *DBH* and *Vigor* were included in the best model (Table 1). *Vigor* reflected the intercept in the *N*_holes_–*DBH* relationship, with the negative value of the coefficient indicating that the intercept increased with decreasing tree vigor. *N*_holes_ of the intact tree (*Vigor* = 5) tended to be approximately twice (=*e*^−0.154×0^/*e*^−0.154×5^) as large as that of dead tree (*Vigor* = 0). Thus, there was a negative relationship between *N*_holes_ and vigor of the white birch.

**Figure 3.**
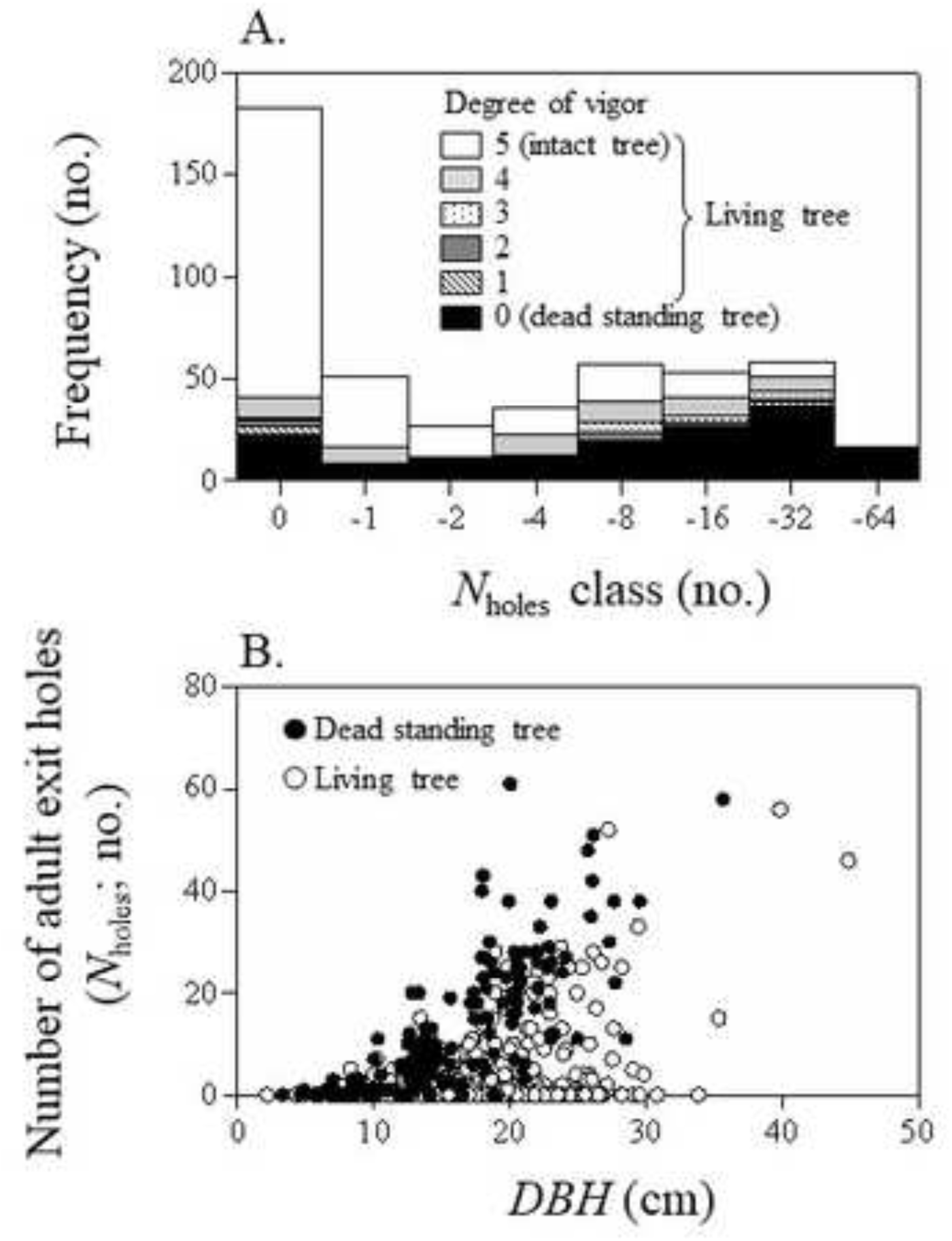
Difference in the number of adult exit holes (*N*_holes_) among living trees and dead standing trees. A) Log-scaled *N*_holes_-class frequency histogram of the number of individuals. B) Relationship between *N*_holes_ and *DBH*, in which living trees are shown by single symbol regardless of degree of vigor.

**Table 1.** Best model for the number of adult exit holes (*N*holes) with respect to *DBH* and tree vigor (*Vigor*). *Coeff.*; estimated coefficient, *Std*.; standard error.

| Variables | *Coeff.* | *Std.* | *z* |
| --- | --- | --- | --- |
| Intercept | -0.582 | 0.496 | -1.174 |
| *DBH* | 0.091 | 0.016 | 5.858 |
| *Vigor* | -0.154 | 0.011 | 14.649 |

Notes: Variance and standard deviation (SD) of the random effect of the intercept and the slope was 2.765 (SD = 1.663) and 0.0013 (SD = 0.036), respectively.

A significant relationship was also found between *N*_holes_ and corresponding dead trees with a minimum *DBH* (*r*^2^ = 0.617, *p* < 0.001; Fig. 4). Thus, *N*_holes_ of small dead trees was less than that of large dead trees. This relationship enables us to calculate a size-dependent threshold for *N*_holes_ for dead or alive of the white birch, i.e., the tree will soon die if *N*_holes_ exceeds the threshold.

**Figure 4.**
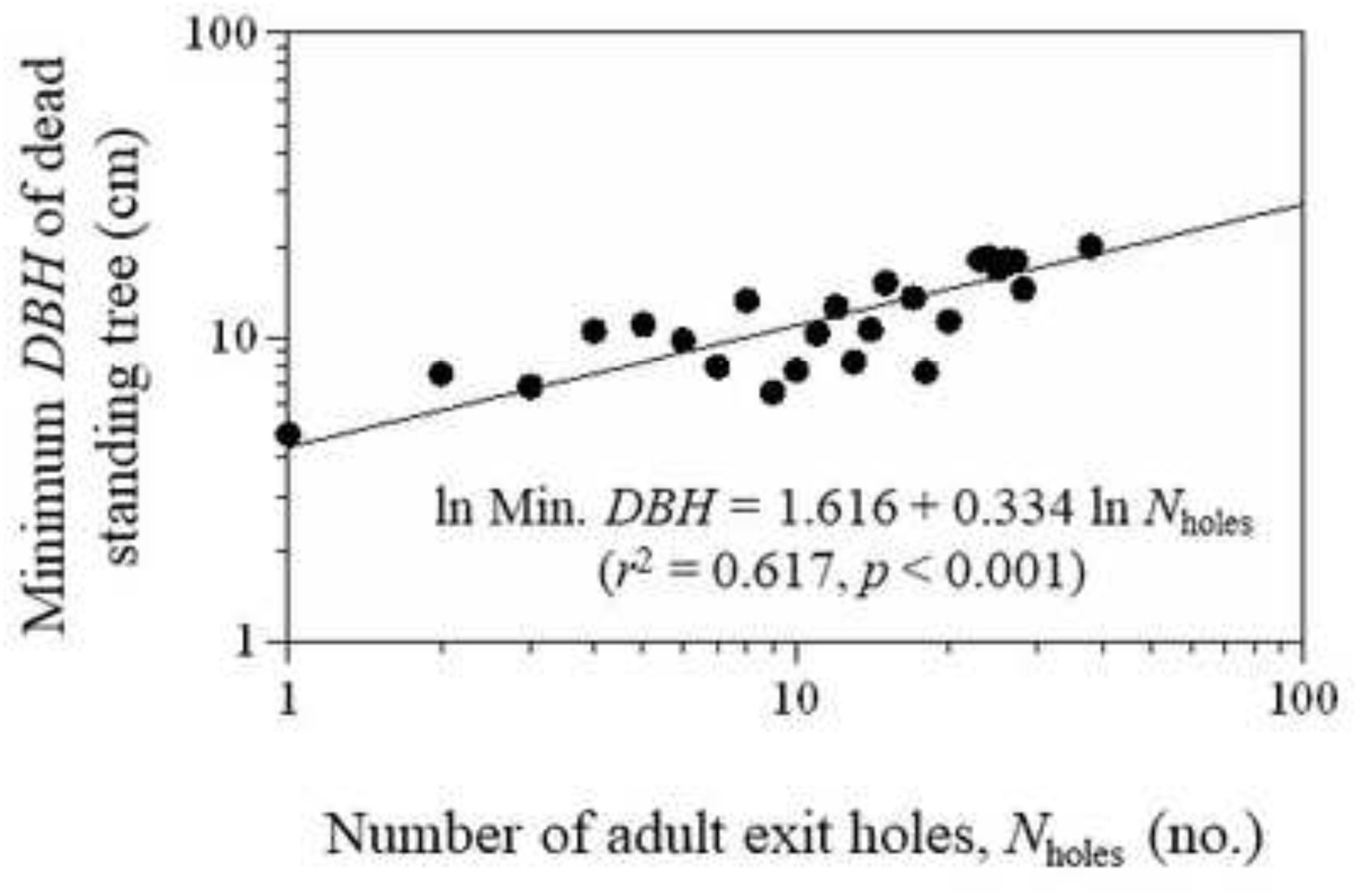
Relationship between the number of adult exit holes (*N*_holes_) and corresponding minimum *DBH* of dead standing trees. We used the data more than 5 individuals for each *N*_holes_ to take the deviation into consideration.

### Diagnosis using RMD

RMD judged that 54.1% (*n* = 33) of the focal trees were diagnosed as *critical,* and 21.3% (*n* = 13) were diagnosed as *suspicious* (Table 2).

**Table 2.** Number of trees for each diagnosis.

| Diagnosis | Number of trees |  |  |
| --- | --- | --- | --- |
|  | Plot 9 | Plot 15 | Total |
| *Not bad* | 13 | 2 | 15(24.6%) |
| *Suspicious* | 9 | 4 | 13(21.3%) |
| *Critical* | 17 | 16 | 33(54.1%) |

Combination 2 (s*uspicious* combines with *critical*) was selected in the best model of GLM, and the interaction term was excluded (Table 3). The intercept of the *N*_holes_–*DBH* relationship varied depending on the severity of diagnosis, whereby *critical* = *suspicious* > *not bad* (Table 3). Since *Diagnosis* gives 0.461 to *critical* and *suspicious*, *N*_holes_ of the trees with *critical* and *suspicious* tended to be 1.586 (= *e*^0.461^/*e*^0^) times greater than that of trees with *not bad*. Hereafter, *suspicious* is combined with *critical*. In Figure 5, estimated *N*_holes_–*DBH* curves of *critical* and *not bad* were layered on the *N*_holes_–*DBH* curves with different vigor indices (Table 1). In both plots, trees with *Vigor* = 1 – 3 would be above *critical* curve. On the other hand, degree of the overlap differed between two plots. Trees with *Vigor* = 4 (long dash line) and 5 (solid line) in Plot 15 would be below *critical* curve, whereas trees with *Vigor* = 5 in Plot 9 would be *critical* to a large extent if *DBH* exceeded ca. 18 cm.

**Figure 5.**
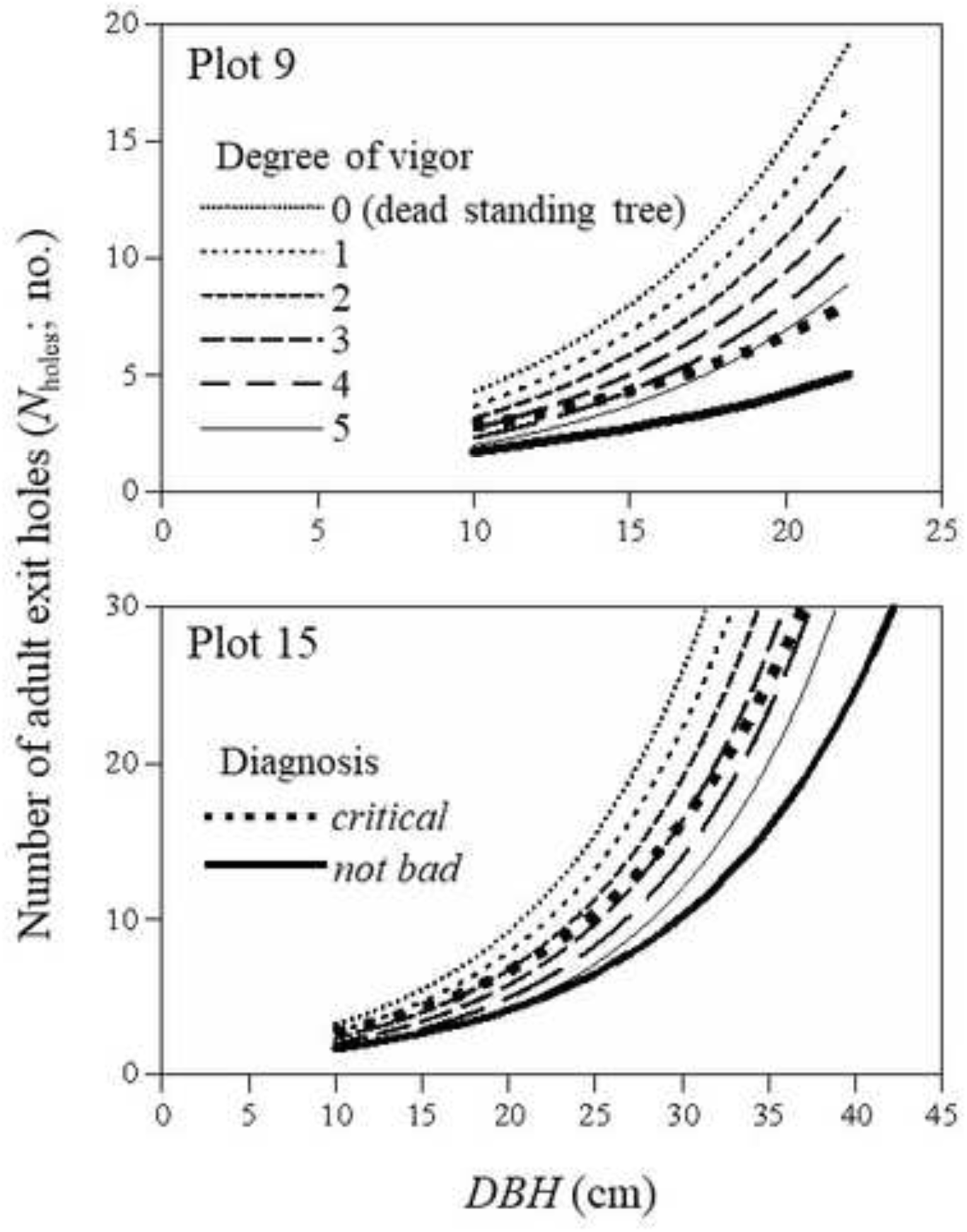
Overlay of diagnosis curve on *N*_holes_–*DBH* curve with different vigor index for Plot 9 (A) and Plot 15 (B). Random effects specific to the plot were calculated by *ranef* function in GLMM analysis. The range of curves correspond to the *DBH* distribution on 14 Apr., 2017; small trees died since Oct., 2015.

**Table 3.** Best model for the number of adult exit holes (*N*holes) with respect to *DBH* and diagnosis. *Coeff.*; estimated coefficient, *Std*.; standard error.

| Variables | *Coeff.* | *Std.* | *z* |
| --- | --- | --- | --- |
| Intercept | -0.334 | 0.160 | -2.088 |
| *DBH* | 0.089 | 0.004 | 19.690 |
| *Diagnosis*: Combination 4 |  |  |  |
| *Suspicious* = *critical* | 0.461 | 0.147 | 3.131 |

Notes: Combination 4: *Suspicious* = *critical*. *Not bad* is reference category (= 0). Null- and residual deviance was 691.42 (df = 60) and 337.08 (df = 58), respectively.

### Probability of the appearance of degraded stands with respect to the severity of infestation

Stand density of the degraded stands tended to be lower than that of intact stands in appearance (Fig. 6). Degraded stands appeared regardless of the year of establishment (Fig. 6).

**Figure 6.**
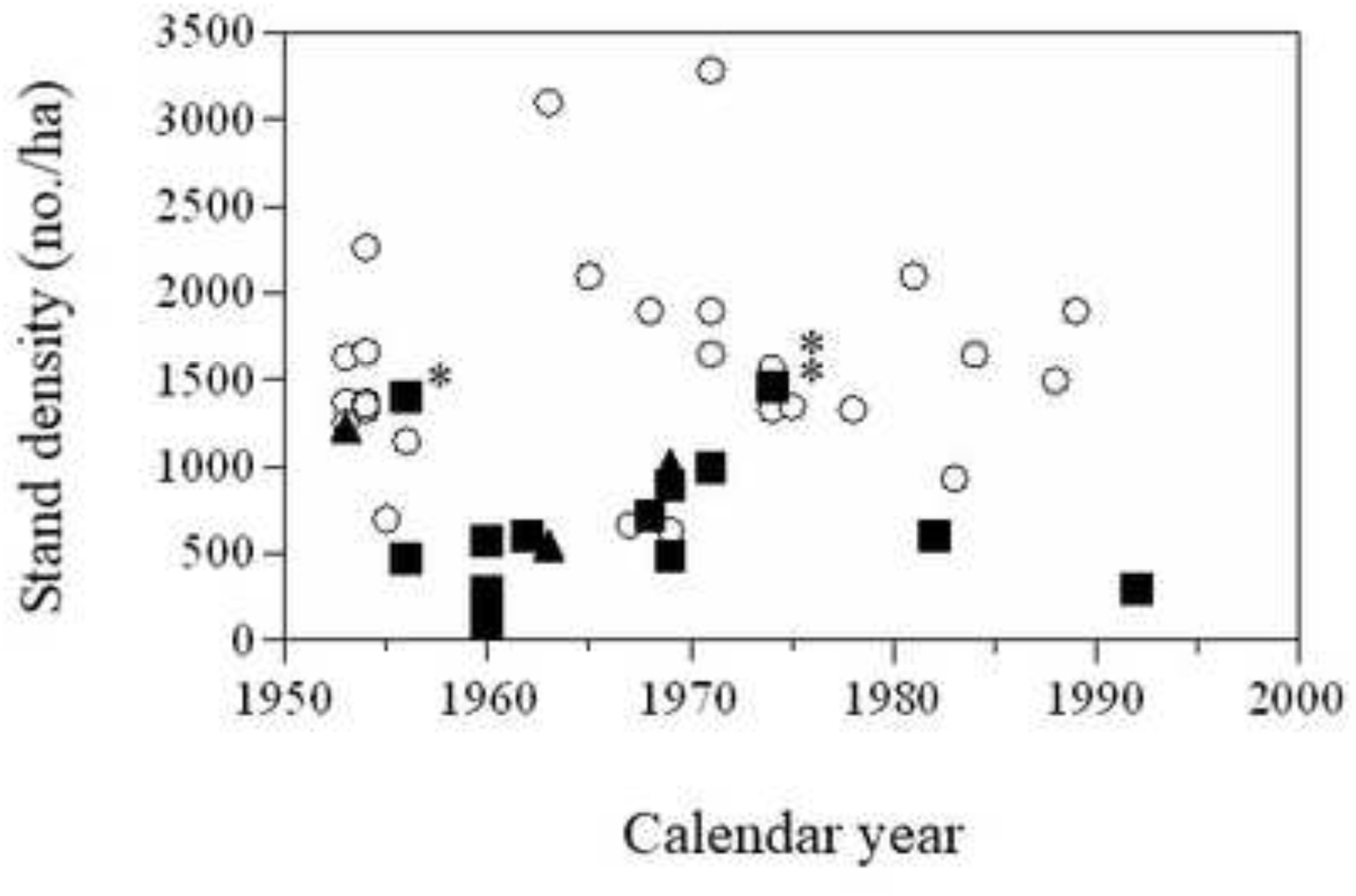
Stand density of each study plot with respect to establish year. ○, intact stands in appearance; ▴, stand just before degradation; ▪, degraded stand. *, Plot 12 was composed of slender trees (see Table S1) that would be associated with relatively high density. ⁑, Plot 21 in which there were many trees with a little foliage in the crown.

The degraded stands appeared, if *N*_D25_ of dead standing tree exceeded 16 (= 2^4^) in the histogram (Fig. 7A; result of GLMM for Eq. 3 was shown in Table 4). If *N*_D25_ exceeded 32 (= 2^5^), almost all stands were degraded. Probability of the appearance of degraded stands with respect to *N*_D25_ could be estimated by following logistic regression model (also see Fig. 7B):

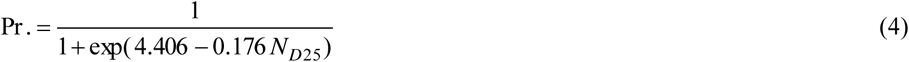

in which AIC of the model was 34.439 (residual deviance (r.d.) = 30.439, degrees of freedom (d.f.) = 49), whereas that of the null model was 65.449. From Eq. 4, 50% of stands would be degraded, if *N*_D25_ would reach 25.0. As a reference, four stands (Plot 9, 32, 38, 41) at a stage just before degradation were shown by different color of bar in Fig. 7A and by different symbols in Fig. 7B.

**Figure 7.**
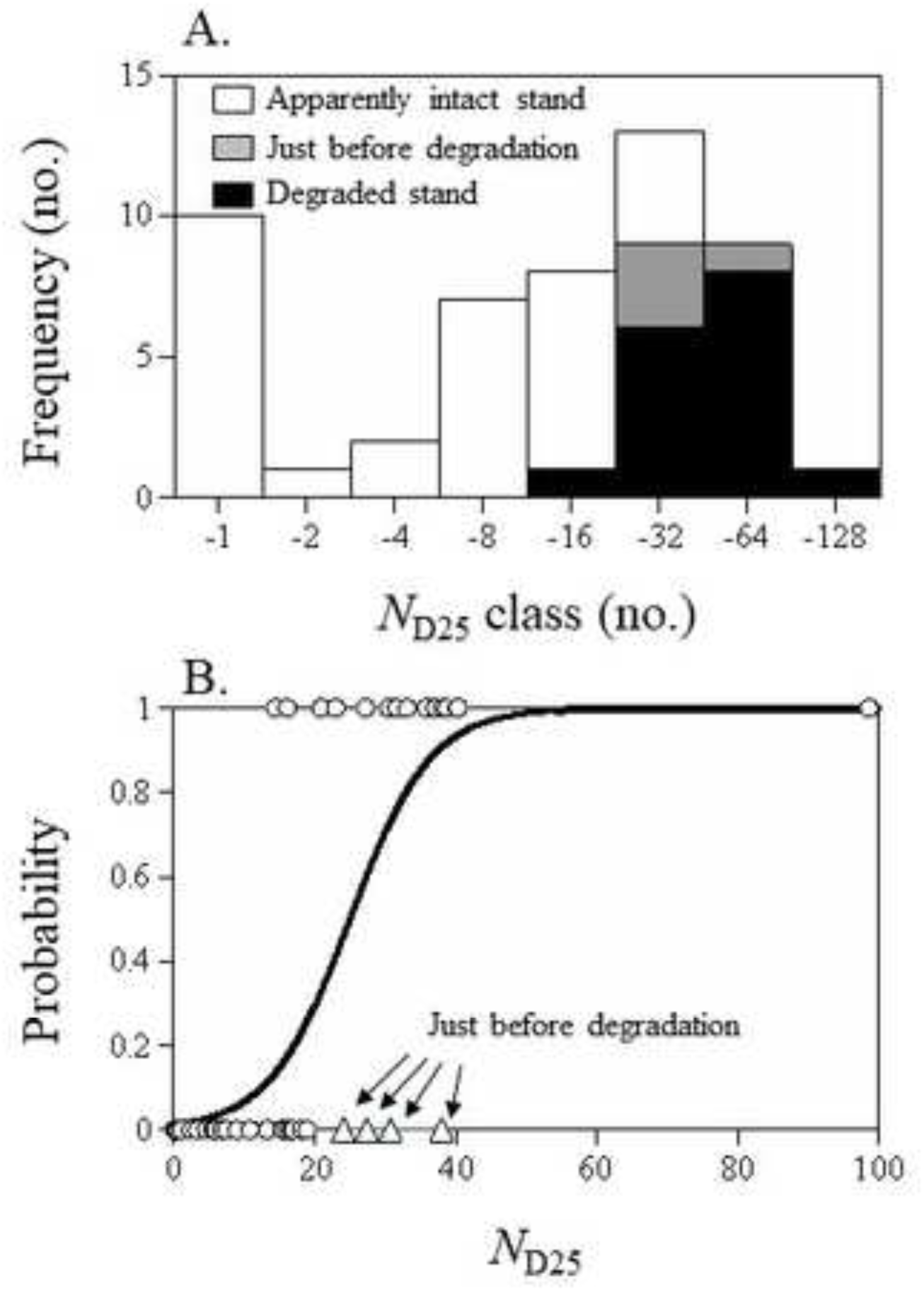
Appearance of degraded stands. A) Log-scaled *N*_D25_-class frequency histogram of the number of stands. B) Logistic regression curve for the probability of the appearance of degraded stand with respect to *N*_D25_.

**Table 4.** Best model for the number of adult exit holes (*Nholes*) with respect to *DBH* and *survival* (*dead* or *alive*). *Coeff.*; estimated coefficient, *Std*.; standard error.

| Variables | *Coeff.* | *Std.* | *z* |
| --- | --- | --- | --- |
| Intercept | -0.973 | 0.285 | -3.411 |
| *Survival_dead* | 0.341 | 0.096 | 3.535 |
| *DBH*×*Survival_alive* | 0.861 | 0.078 | 11.074 |
| *DBH*×*Survival_dead* | 1.034 | 0.084 | 12.333 |

Notes: *Survival\_alive* is the reference category (= 0). Variance and standard deviation (SD) of the random effect of the intercept and the slope was 3.384 (SD = 1.840) and 0.0158 (SD = 0.398), respectively.

## Discussion

The number of adult exit holes of the white-spotted longicorn beetle (*N*_holes_) in dead Japanese white birch trees was greater than that in living trees (Fig. 3), and the degradation of stands tended to progress with increasing *N*_holes_ (Figs. 7a, b). These facts imply that *N*_holes_ is a useful and simple index to explain the severity of infestation by the white-spotted longicorn beetle. There were certainly dead trees without adult exit holes (Fig. 3A), especially small trees. Generally, mortality in small trees was caused by shading. As white birch is a pioneer species, suppression often causes the mortality of small trees in this species.

Larvae of the white-spotted longicorn beetle bore tunnels in the cambial region and sapwood (Figs. 1D and F; cf. Haack et al. 2010). After repeated attack, larval galleries disrupt the tree’s vascular tissues, and it can lead to tree death (cf. Haack et al. 2010). Thereafter, the dieback of crown could reflect the severity of infestation in relation to the degree of disruption of the cambial region. It should be noted that we may have to pay attention to the possibility that some larvae that do not pupate and develop into adults also bore tunnels in the wood (cf. Fig. S3B) and, therefore, do not create exit holes (cf. Adachi 1989). However, at the moment, there is no information about the mortality of the larvae on Japanese white birch. For example, Golec et al. (2018) investigated the mortality of Asian longhorned beetle on willows and popular in China, and reported that on average 59.3% of eggs were killed by undetermined factors. On average 8.3%, 15.6% and 9.1% of immatures (larvae and pupae) were killed by undetermined factors, woodpeckers, and unidentified predators, respectively. 10.1% and 8.1% of adults that did not emerge were killed by undetermined factors and parasites, respectively. It implies that the number of adult exit holes surveyed in this study underestimates the severity of infestation.

The tolerance of trees to the infestation must be size dependent, since the cambial region is proportional to *DBH* to a large extent. This implies that the maximum cumulative capacity of larvae in each tree is also size dependent. Adachi (1989) also found the positive relationship between the number of adult exit holes of white-spotted longicorn beetle and tree diameter in citrus. The tendency implies that the lethal threshold in *N*_holes_ is size-dependent. We may be able to use it as simple diagnostic criteria for on-site decisions, i.e., the regression line shows the critical stage of the infestation. If *N*_holes_ of the tree would reach the line, the tree would most likely die soon.

RMD examination revealed that many trees in apparently intact stands were already in a critical condition due to many exit holes, especially in Plot 9 (Fig. 5 and Table 1). Since Plot 9 and Plot 15 are close to degraded stands (Fig. 2), it is not a surprise that these stands will soon degrade. In both plots, trees with *Vigor* = 1 – 3 would be above *critical* curve (Fig. 5) implying that the wood decayed severely (cf. Fig. S1). These trees will soon die. Results of the RMD examination will reinforce the diagnosis using simple diagnostic criteria based on the relationship between *N*_holes_ and the corresponding minimum *DBH* in dead trees (Fig. 4). Thus, the RMD examination evaluates the risk of tree falling due to the wood decay, whereas the simple diagnostic criteria evaluate the risk of tree mortality. On the other hand, degree of overlap between *critical* curve and curve of trees with *Vigor* = 5 differed among two plots (Fig. 5). It is considered that the difference was caused by the difference in progress of infestation; i.e., *N*_D25_ of dead standing tree was 27.4 in Plot 9 and 18.1 in Plot 15, respectively (Table S1). It means that the infestation was progressing in Plot 9 more than Plot 15. Even if the appearance of crown looks intact, it does not necessarily guarantee the tree health. Disease is steadily progressing inside. Unless the foliage declines excessively, the number of adult exit holes will prior to the crown vigor for visual inspection.

The year of establishment was not strongly correlated with degradation (Fig. 6A) implying that the degradation had occurred recently. It is currently unclear why the mass mortality of the white birch shelterbelts caused by the white-spotted longicorn beetle occurred in central Hokkaido. For example, the Asian longhorned beetle is originally from China, and is a harmful invasive species in Europe and North America (Favaro et al. 2015; Haack et al. 2010). However, both white-spotted longicorn beetle and Japanese white birch are native species in Hokkaido. It does not matter whether the pest is native or non-native in our study. According to Arango-Velez et al. (2013), on the other hand, stressed trees are more susceptible to attack by insects than their healthy counterparts. Since Japanese white birch shows little tolerance to flooding (Terazawa and Kikuzawa 1994), the trees in the shelterbelts might lose vigor with age due to relatively high ground water level in the drained peatland. It may explain the relatively high susceptibility to the white-spotted longicorn beetle. Monoculture might also accelerate the mass mortality.

Severely infested stands will eventually lead to degradation. Degraded stand will appear, if *N*_D25_ exceeds 16 (Fig. 7A). The logistic regression curve implies that 50% of stands are degraded if *N*_D25_ exceeds 25 (Fig. 7B). These results mean that the destruction of the white birch stands caused by the white-spotted longicorn beetle proceeded rapidly. It is no wonder that there are many adult white-spotted longicorn beetles in the stands in which degradation is progressing. Trees should be removed as earlier as possible, if *N*_D25_ exceeds 16. The infested wood must be burned or fumigated after cutting to inhibit adult emergence any more, that is, there will be pupae in the wood yet.

No effective pest control technique has been developed for *Anoplophora* spp. on any of their hosts (Haack et al. 2010). Because the larvae of the white-spotted longicorn beetle bore into the trunks, they are difficult to control and damage to the hosts is often difficult to detect (Fujiwara-Tujii et al. 2016). Haack et al. (2010) pointed out the effectiveness of biological control using fiber bands containing the entomopathogenic fungi, *Beauveria brongniartii* and *Metarhizium anisopliae*. This method may be effective for road side trees in urban area, whereas it seems unrealistic for shelterbelts and natural forests because of huge number of trees. Though Japanese white birch can regenerate by stump sprouting (Masaka et al. 2000), this is not possible if the roots are rotting. It means that natural recovery of the degraded shelterbelts is not possible. Therefore, we may have no choice but to exchange this tree species for an alternative shelterbelt species in central Hokkaido.

As the degraded shelterbelts were scattered sporadically throughout our study area (Fig. 2), it seems that the white-spotted longicorn beetle attacks can occur anywhere. Indeed, one degraded road-side tree line composed of the Japanese white birch was found in Sapporo in 2017, about 50 km from Bibai, in the census of road side trees by Sapporo City Hall (Sapporo City Hall, unpublished data). If the infestation area would expand for the future, we might be asked to set priorities to remove the infested stands. Visual inspections based on the number of adult exit holes (Figs. 4 and 7A, B) together with RMD examination (Fig. 5) will contribute to the diagnosis of infested white birch.

## Acknowledgments

We are grateful to Sorachi General Subprefectural Bureau and Bibai City Hall for permission to cut the trees in the regally protected wind shelterbelt; to Abe T., Kobayashi Y., Kokubo A., Hayasaka K., Nagasawa R., Nishimura S., Tanahashi I. at HFRI for their help in field investigation (in alphabetical order).

*Anoplophora* spp. is the most damaging wood-boring pests in the Northern Hemisphere. The aim of our study is to develop the diagnosis method for decline of white birch tree using the number of adult exit holes of the white-spotted longicorn beetle. In addition to the visual inspection, we also applied the decay diagnosing method of wood using the resonance measurement device (RMD) in corroboration of the effect of infestation on the wood. Tree vigor was negatively influenced by the number of adult exit holes, and we found a size-dependent lethal threshold in the number of adult exit holes. RMD examination revealed that many trees in apparently intact stands were already in a critical condition due to many exit holes. Even if the appearance of crown looks intact, it does not necessarily guarantee the tree health. Unless the foliage declines excessively, the number of adult exit holes will prior to the crown vigor for visual inspection. We also demonstrated the appearance of degraded stands with subject to the number of adult exit holes. Visual inspections based on the number of adult exit holes together with RMD examination will contribute to the diagnosis of infested white birch.

---

**Figure S1.**
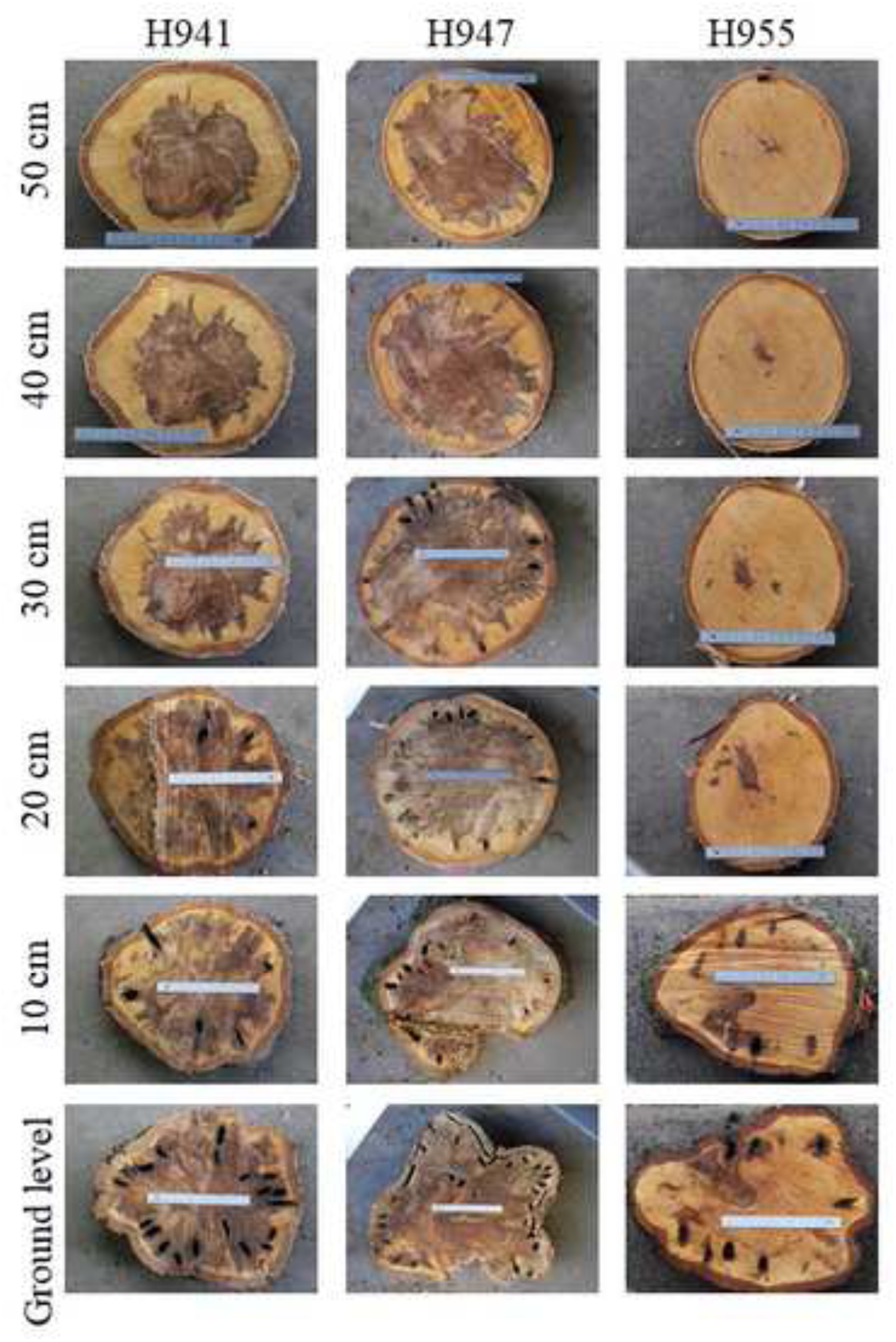
Examples of cross section of infested trees (H941, H947 and H955) with different height above the ground. These trees were used for calibration of RMD diagnosis. Wood decayed severely in H941 and H947, while wood was almost intact in H955. Scale = 15 cm.

**Figure S2.**
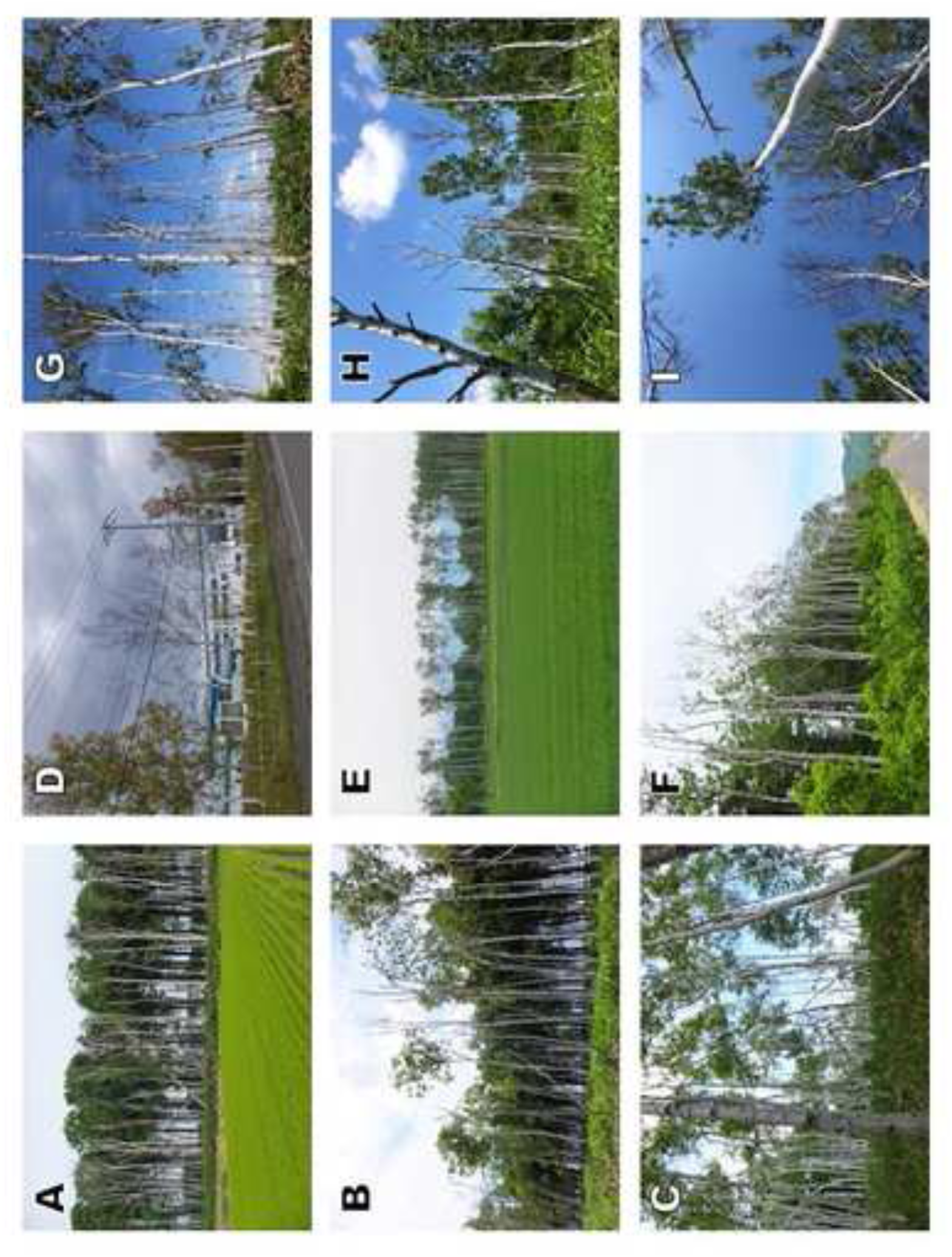
Focal degraded stands in Iwamizawa, Mikasa, Shinshinotsu, and Tsukigata. A) Plot 24 (15 Jun. 2017), B) Plot 25 (16 Jun. 2017), C) Plot 26 (16 Jun. 2017), D) Plot 29 (16 Oct. 2016), E) Plot 30 (7 Jun. 2017), F) Plot 31 (7 Jun. 2017), G) Plot 37 (16 Jun. 2017), H) Plot 46 (13 Jun. 2017), I) Plot 51 (14 Jun. 2017). Notes that we have no photos about Plot 21 in growing season because the investigation was carried out on 29 Mar., 2017. Crown vigor in Plot 21 was observed during the growing season in 2016. In A, B, C and F, shelterbelt is composed of two belts; one is Norway spruce and the other is white birch. The belt of Norway spruce is found behind the belt of white birch except C.

**Figure S3.**
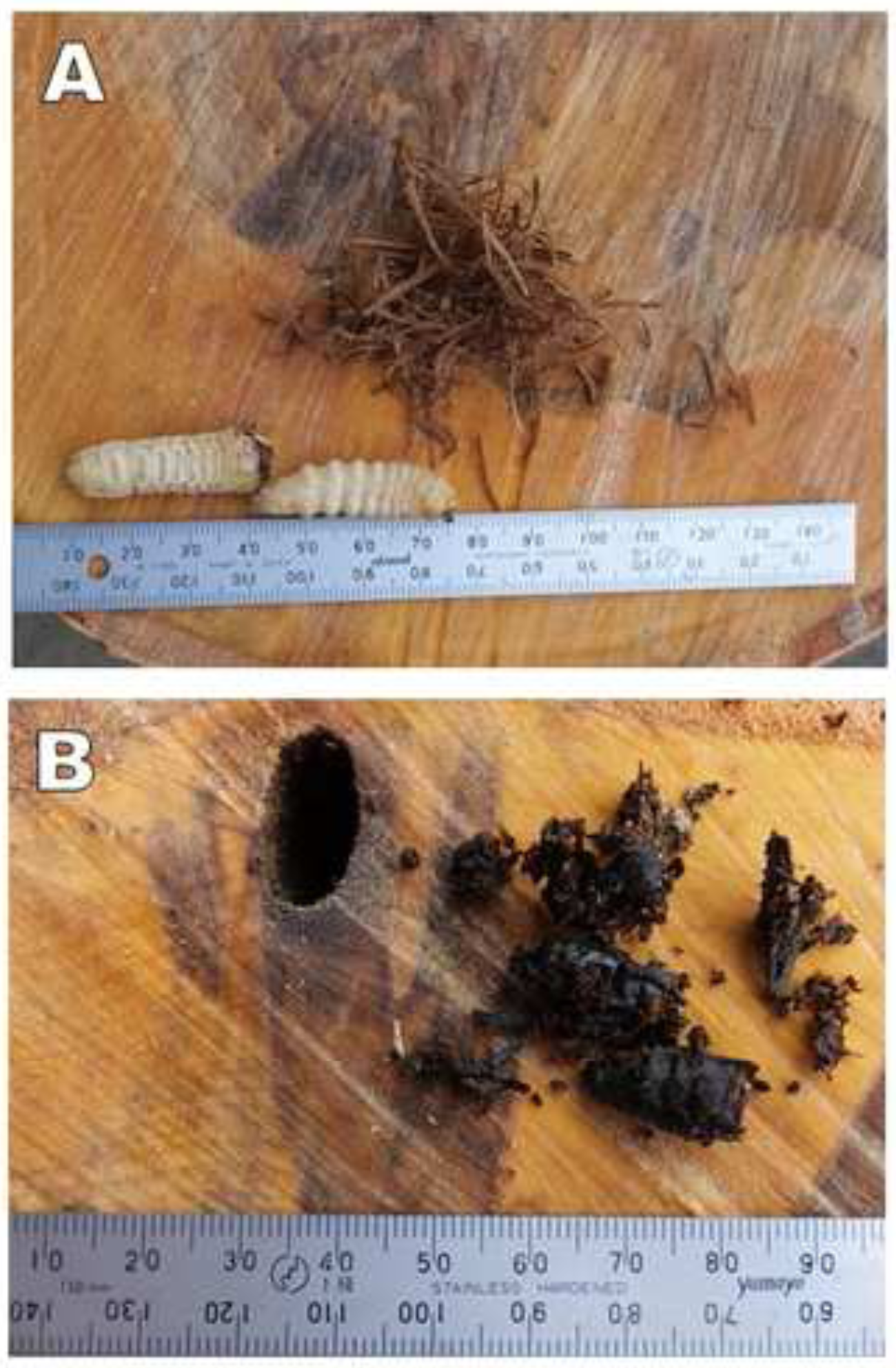
Stuffing in the tunnel bored by larvae of white-spotted longicorn beetle. A) Larvae and sawdust. B) Rotten and blackening sawdust and adult dead body.

**Table S1.**
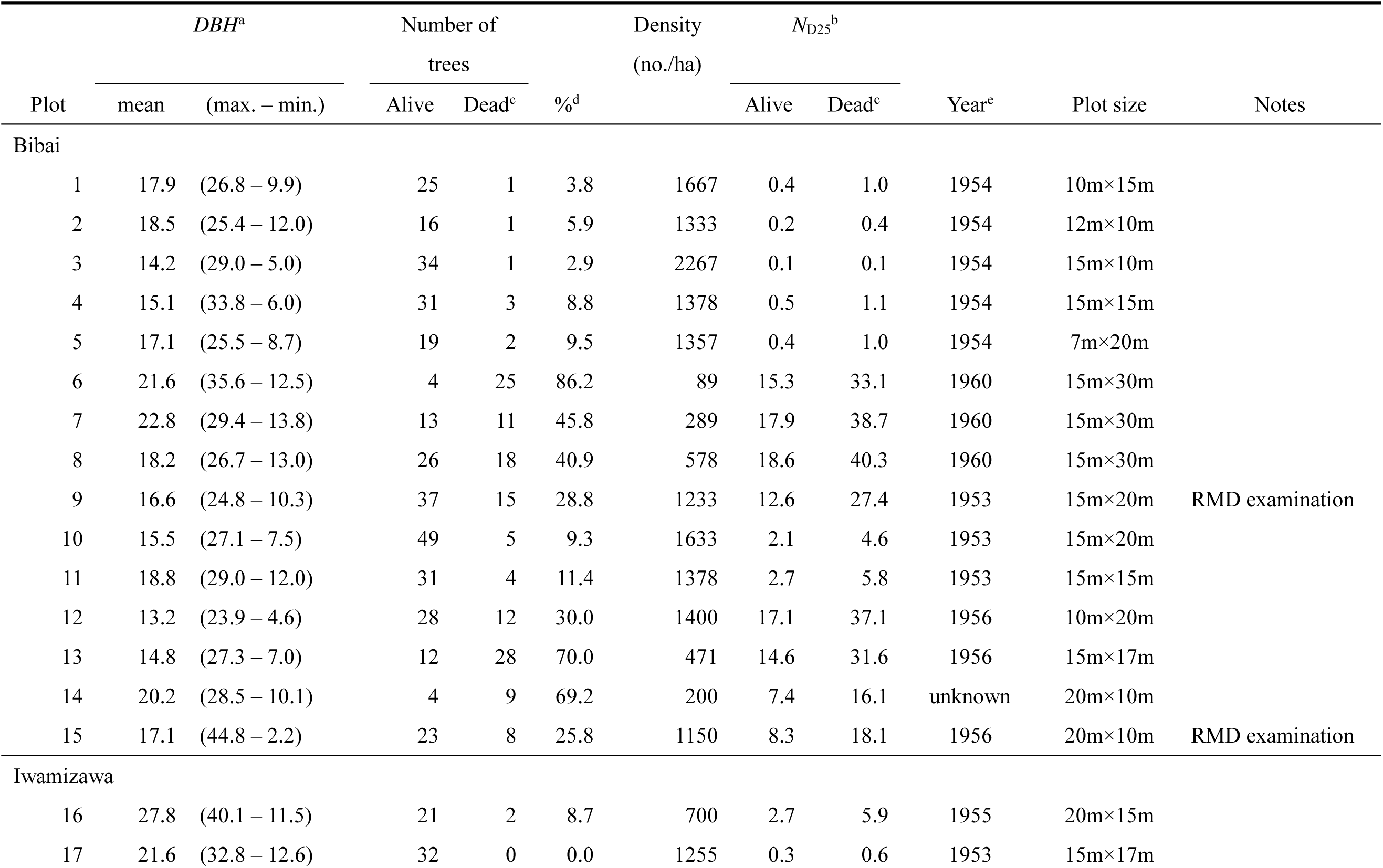
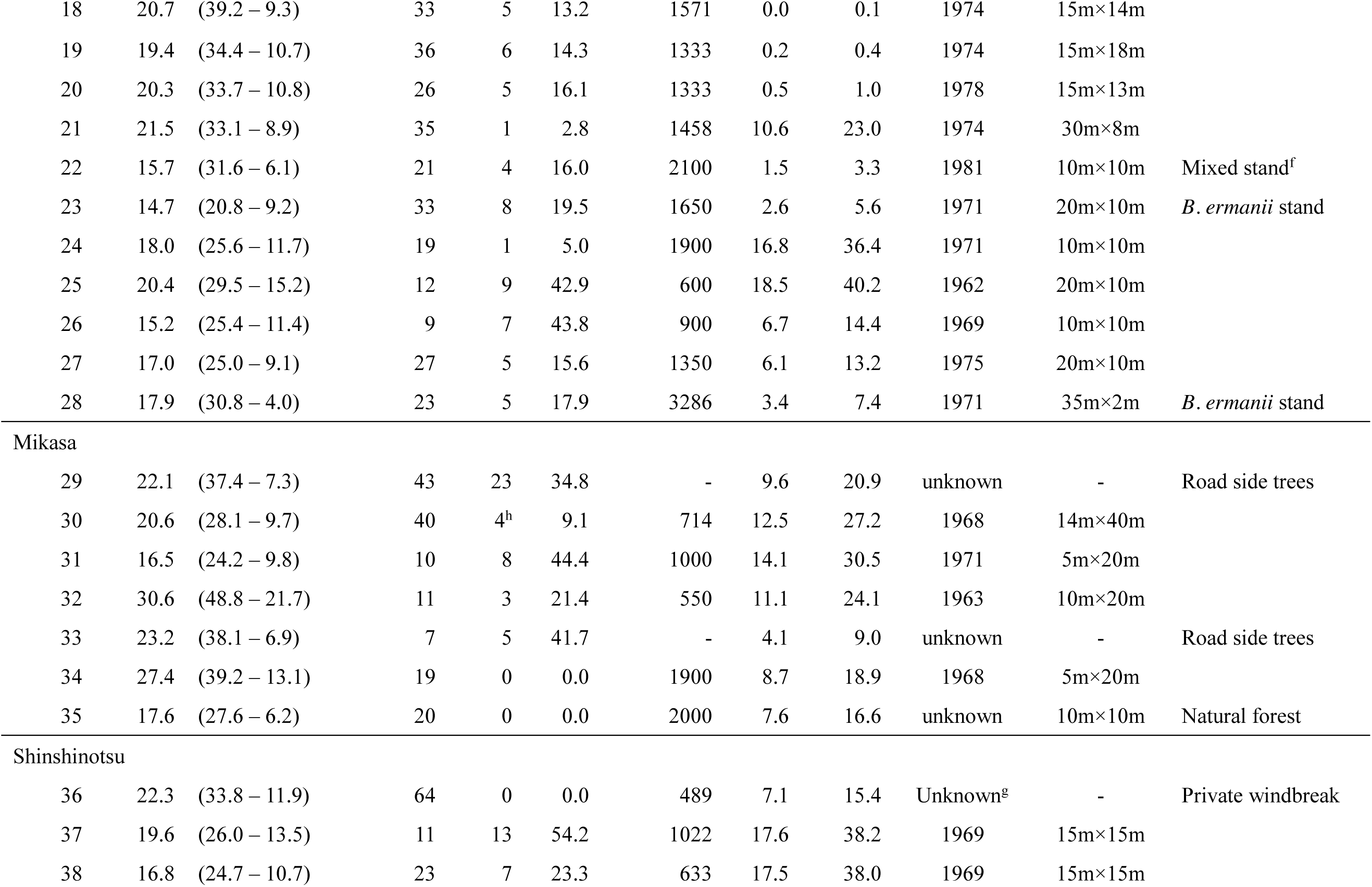
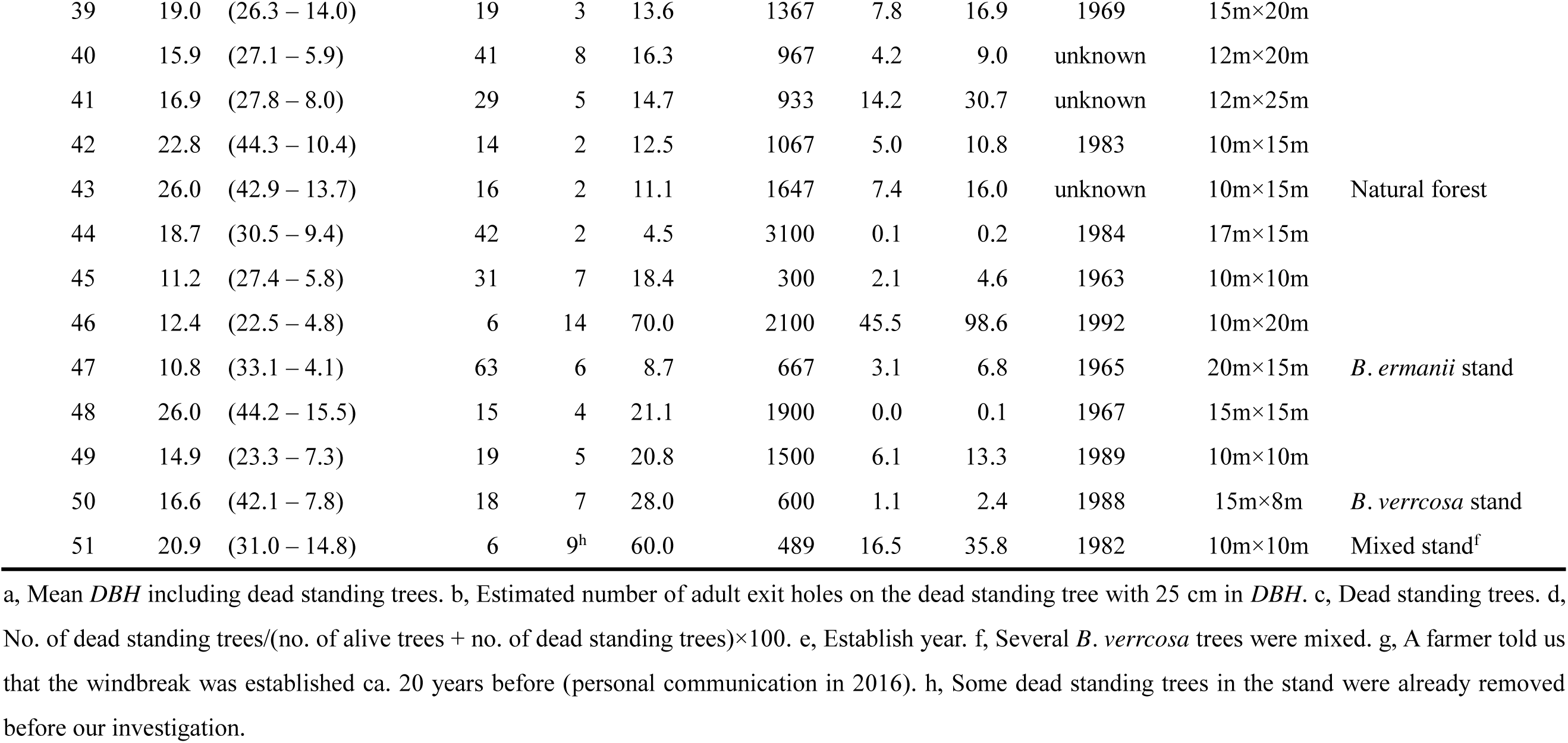
Stand characteristics.

